# Proteomic associations with cognitive variability as measured by the Wisconsin Card Sorting Test in a healthy Thai population: A machine learning approach

**DOI:** 10.1101/2024.10.25.620177

**Authors:** Chen Chen, Bupachad Khanthiyong, Benjamard Thaweetee-Sukjai, Sawanya Charoenlappanit, Sittiruk Roytrakul, Phrutthinun Surit, Ittipon Phoungpetchara, Samur Thanoi, Gavin P Reynolds, Sutisa Nudmamud-Thanoi

## Abstract

Cognitive function is the term for the higher-order mental processes in the brain that gather and process information, and it mirrors brain activity. Cognitive function in adults exhibits variability as a result of genetic and environmental components such as gender, age[1], and lifestyle factors to name a few. Interindividual variability in cognitive trajectories has been observed in community-dwelling older adults across different cognitive domains. Inter-individual variations in cognitive response to identical physical exercise are also evident. This study aimed to explore the association between serum protein expression profiles and one measure of cognitive variability, as measured by the Wisconsin Card Sorting Test (WCST), in a healthy Thai population using a machine learning approach. This study included 199 healthy Thai subjects, ranging in age from 20 to 70 years. Cognitive performance was measured by the WCST, and the WCST % Errors was used to define the lower and higher cognitive ability groups. Serum protein expression profiles were studied by the label-free proteomics method. The Linear Model for Microarray Data (LIMMA) approach in R was utilized to assess differentially expressed proteins (DEPs) between groups; subsequently bioinformatic analysis was performed for the functional enrichment and interaction network analysis of DEPs. A random forest model was built to classify subjects from the lower and higher cognitive ability groups. Cross-validation was used for model performance evaluation. The results showed that, there were 213 DEPs identified between the poor and higher cognition groups, with 155 DEPs being upregulated in the poor cognition group. Those DEPs were significantly enriched in the IL-17 signaling pathway. Furthermore, the analysis of protein-protein interaction (PPI) network revealed that most of the selected DEPs were linked to neuroinflammation-related cognitive impairment. The random forest model achieved a test classification accuracy of 81.5%. The model’s sensitivity (true positive rate) was estimated to be 65%, and the specificity (true negative rate) was 85.9%. The AUC (0.79) indicates good binary classification performance. The results suggested that a measure of poor WCST performance in healthy Thai subjects might be attributed to higher levels of neuroinflammation.

## Introduction

Cognitive function is the term for the higher-order mental processes in the brain that gather and process information, and it mirrors brain activity. Cognitive function in adults exhibits variability as a result of genetic and environmental components [2, 3], such as gender, age [1], and lifestyle factors [3], to name a few. Interindividual variability in cognitive trajectories has been observed in community-dwelling older adults across different cognitive domains [4]. Inter-individual variations in cognitive response to identical physical exercise are also evident [5, 6].

Furthermore, these factors may have complex direct and indirect interactions on normal cognitive variability. Self-rated health, for example, has been proposed as a mediator variable in the relationship between life satisfaction and physical and mental health among the elderly in Brazil [7]. Our prior studies have suggested that the cognitive sex differences in WCST performances between healthy Thai males and females may be a result of their susceptibility to the N-methyl-D-aspartate receptor (NMDAR)-mediated excitotoxicity [8], and that the interaction between the cholinergic system and estrogen might contribute to the effect of schooling in attenuating such sex-dependent cognitive differences [9]. As a result, investigating the complex interactions between such factors using information technology and computer-based algorithms might enhance our understanding of the variability in normal cognition.

Machine learning algorithms have been employed to classify cognitive profiles [10–12] as well as predict cognitive health in the global population [13–15]. When using machine learning methods, the racial and ethnic background of subjects is an essential issue to take into account [16], since racial bias is a prevalent challenge facing machine learning in human studies [17]. Because cognitive function is a reflection of brain activity, alterations in molecular factors such as neuronal and neurotransmission-related proteins may provide additional information for understanding cognitive variability in healthy people. Therefore, this study aimed to explore the association between serum protein expression profiles and one measure of cognitive variability, as assessed by the Wisconsin Card Sorting Test (WCST), in a healthy Thai population by employing a machine learning approach.

## Methods

### Participants and data

This study included 199 healthy Thai subjects, ranging in age from 20 to 70 years, with samples collected and cognitive tests conducted from October 20, 2014, to August 25, 2018, as previously reported [18]. The researchers assessed the archived samples and data from July 6, 2022, to July 6, 2023, for the preparation of this publication; all subjects were assigned a study ID that concealed individual identities. The cognitive performance of each participant was evaluated using the WCST, a test that measures cognitive flexibility [19, 20]—the capacity to adapt the behavioral response mode to changing conditions—which is commonly employed to assess frontal lobe function, especially the prefrontal cortex [21]. 3 ml of blood was collected from the cubital vein of each subject immediately following the completion of the WCST. The serum protein expression profiles of the subjects were then analyzed using the label-free proteomics method, as previously described [8].

The mean and standard deviation (SD) of the percentage of total errors (%Errors) in WCST were computed in male and female subjects separately, covarying for age [8]. A negative correlation has been observed between WCST %Errors and the Full-Scale Intelligence Quotient (FSIQ) [22, 23], a metric used to assess a person’s overall level of general cognitive and intellectual functioning [24]. Male and female subjects with %Errors > 1SD above their sex-specific mean were considered to have poor cognitive performance and were assigned to the lower cognitive ability group [25]. All methods were performed in compliance with relevant guidelines and regulations (Declaration of Helsinki). The Institutional Review Board (IRB) of Naresuan University, Thailand approved this research (COA No. 0262/2022). Participation was voluntary, and written informed consent was obtained from each subject.

### Bioinformatic analysis

Differentially expressed proteins (DEPs) between the two cognition groups were identified using the Linear Model for Microarray Data (LIMMA) approach in R version 4.2.3 [26], with P≤0.01 considered significant.

Pathway analysis and Gene Ontology (GO) enrichment analysis were performed on the Database for Annotation, Visualization, and Integrated Discovery (DAVID) [27, 28]. The protein-protein interaction network was studied using Pathway Studio version 12.5 [29]. DEP expression levels were displayed by Multi-Experiment Viewer (MeV) version 4.9.0 [30]. P≤0.05 was considered significant.

### Preprocessing

The training and testing datasets were respectively proportioned at 0.6 and 0.4 of the total sample. To address the small proportion of subjects defined as having poor cognitive performances, the synthetic minority over-sampling technique (SMOTE) was employed [31]. In contrast to up sampling, which involves the replication of duplicate samples, SOMTE generates synthetic data that closely resembles the initial data set. SMOTE was exclusively applied to the training dataset during cross-validation to minimize overfitting. The testing dataset, which was not included in SMOTE or cross-validation procedures, was used to evaluate the final performance metrics of the model [13].

The DEPs that were significantly enriched in cognition-related pathways, as identified by GO enrichment analysis, pathway analysis, and protein-protein interaction network analysis, were selected as model variables. Age was also included in the model since it is strongly connected to cognitive impairment in the healthy population [32, 33].

### Machine learning model

The machine learning (ML) processes were performed using *tidy models* in R [34]. Tidy model is a meta-package for modeling and statistical analysis. It allows for the creation of a unitary preprocessing dataset, ensuring a reliable comparison across various ML models, with the random forest model being employed here. The principles and procedures of the random forest model are thoroughly discussed in multiple publications [35–37]. In brief, random forest is a decision tree algorithm-based ensemble learning method applicable to both classification and regression analysis [38]. Typically, approximately two-thirds of the study samples are allocated to model fitting or training, with the other one-third applied for model testing, known as out-of-bag (OOB) data, which is employed for evaluating the performance of the model. When it comes to classification, the OOB error rate—the rate at which OOB samples are misclassified over various classes—can be used to quantify how well the model is performing. The significance of each variable in the final model is assessed using a ranked measure of variable importance.

### Model validation

Cross-validation is a recommended evaluation method used in machine learning to determine how well the models work [39]. In this study, a 10-fold cross-validation was conducted on the training dataset to evaluate the models’ performances.

In addition, model hyperparameters were tuned during the cross-validation [34]. Specifically, the following hyperparameters were tuned in this study: the number of trees was 1000, the minimum number of data points required for a node further splitting was 2, and 10 predictors were randomly sampled at each split when building the tree models.

### Performance metrics

In this study, the overall accuracy, Matthew’s correlation coefficients (MCC), F_1_-score, and area under the receiver operating characteristic curve (AUC) were used as performance metrics for the random forest model [13, 14].

## Results

### Demographic data of the study population

The demographic characteristics of the study population are detailed in Table 1. The mean age of the participants was 45.6±19.3 years, with 55.3% being female. The cutoff point as defined by %Errors > 1SD from the mean in males was 62.3, whereas in females, it was 67.4.

**Table 1.** Demographic data of the study population.

|  | Lower cognitive ability (n=41) | Higher cognitive ability (n=158) |
| --- | --- | --- |
| Age, years | 51.5±17.5 | 44.0±19.6 |
| Sex, female | 22 (53.7%) | 88 (55.7%) |
| Education, secondary & tertiary | 14 (34.1%) | 93 (58.9%) |
| WCST %Errors | 69.6±4.78 | 48.6±10.0 |

Data is shown as mean±SD

### Bioinformatic analysis

There were 213 differentially expressed proteins identified between the poor and higher cognition groups, with 155 DEPs being upregulated in the poor cognition group and the remaining 58 DEPs being downregulated (see Fig 1). Data from the subsequent GO enrichment analysis demonstrated that DEPs were substantially enriched in the following biological pathways: regulation of protein stability (P=0.01), macromolecule methylation (P=0.03), regulation of neuron projection development (P=0.04), and retinoic acid catabolic process (P=0.04), as shown in Table 2. Regarding the KEGG pathway analysis [40–42], these DEPs showed a significant enrichment in the IL-17 signaling pathway (P=0.05), see Table 3.

**Fig. 1.**
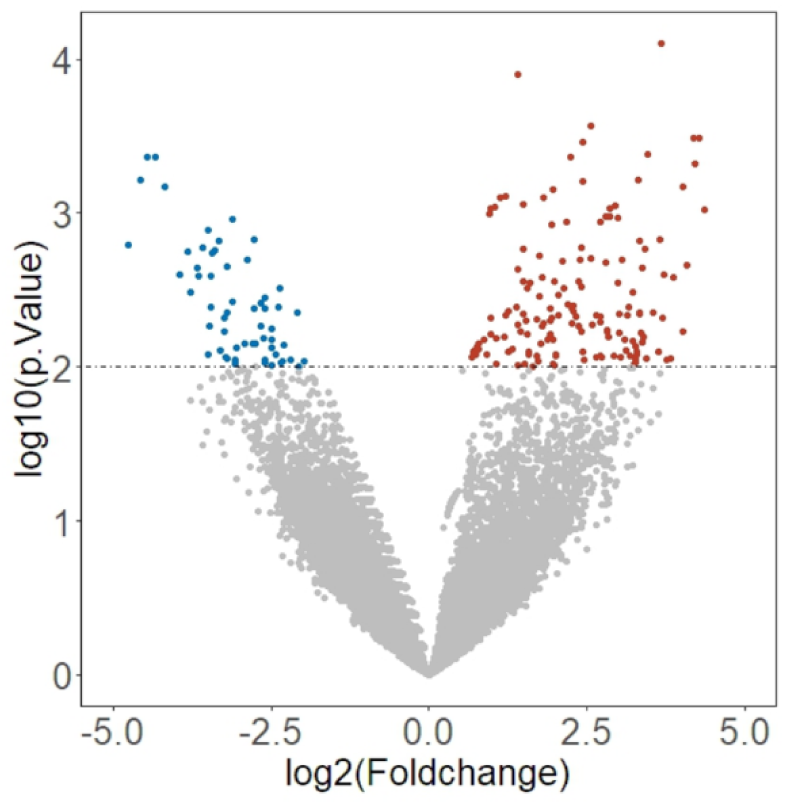
Volcano plot of DEPs between poor and normal cognition groups. Color blue represents DEPs that are downregulated in the poor cognition group, red represent DEPs unregulated in poor cognition group.

**Table 2.** Gene Ontology analysis of DEPs between lower and higher cognitive ability groups.

| Number | Pathway ID | Pathway | Mapped DEPs | Fold<br>Enrichment | P-value |
| --- | --- | --- | --- | --- | --- |
| 1 | GO:0031647 | Regulation of protein stability | GET4, CCAR2,<br>KAT2A, UPS25,<br>INSC | 5.84 | 0.01 |
| 2 | GO:0043414 | Macromolecule methylation | FTSJ3, METTL4 | 55.5 | 0.03 |
| 3 | GO:0010975 | Regulation of neuron projection<br>development | NCS1, BRSK2,<br>NTNG1 | 8.99 | 0.04 |
| 4 | GO:0034653 | Retinoic acid catabolic process | CRABP1, CYP2W1 | 44.3 | 0.04 |

**Table 3.**
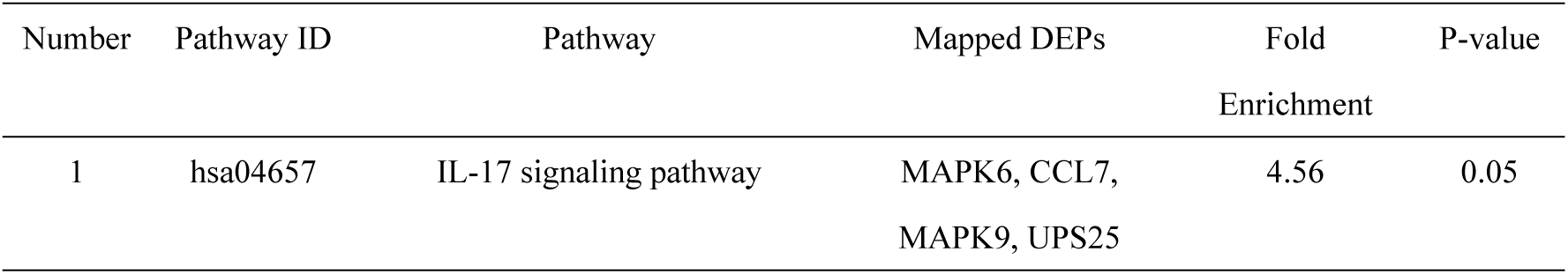
Pathway analysis of DEPs between lower and higher cognitive ability groups.

Furthermore, protein-protein interaction (PPI) network analysis indicated that most of the 16 DEPs enriched in the aforementioned pathways were linked to neuroinflammation-related cognitive impairment. The other four DEPs found to be involved in this PPI network were: serotonin receptor 7 (HTR7), metabotropic glutamate receptor 4 (GRM4), choline transporter like protein 2 (SLC44A2), and pro-adrenomedullin (ADM) (see Fig 2). Glutamatergic [43], serotonergic [44], and cholinergic [45] systems have been shown to have a role in cognitive impairment. Additionally, ADM has been suggested as a potential biomarker for cognitive impairment [46, 47]. As a result, all those 20 proteins were included as variables in the random forest model.

**Fig. 2.**
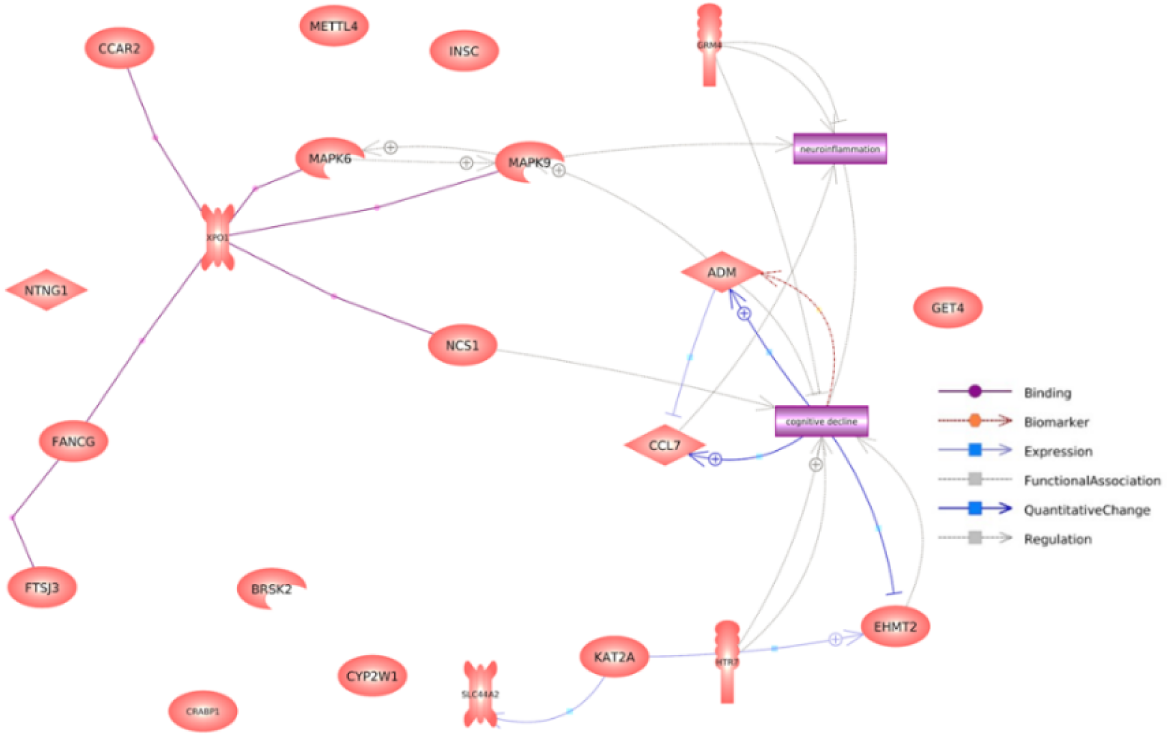
Protein-protein interaction network of selected DEPs

### Model performance

The random forest model achieved a test classification accuracy of 81.5% (see Table 4). The model’s sensitivity (true positive rate) was estimated to be 65%, while the specificity (true negative rate) was 85.9%. The AUC (0.79) indicates good binary classification performance [48] (see Fig 3).

**Fig. 3.**
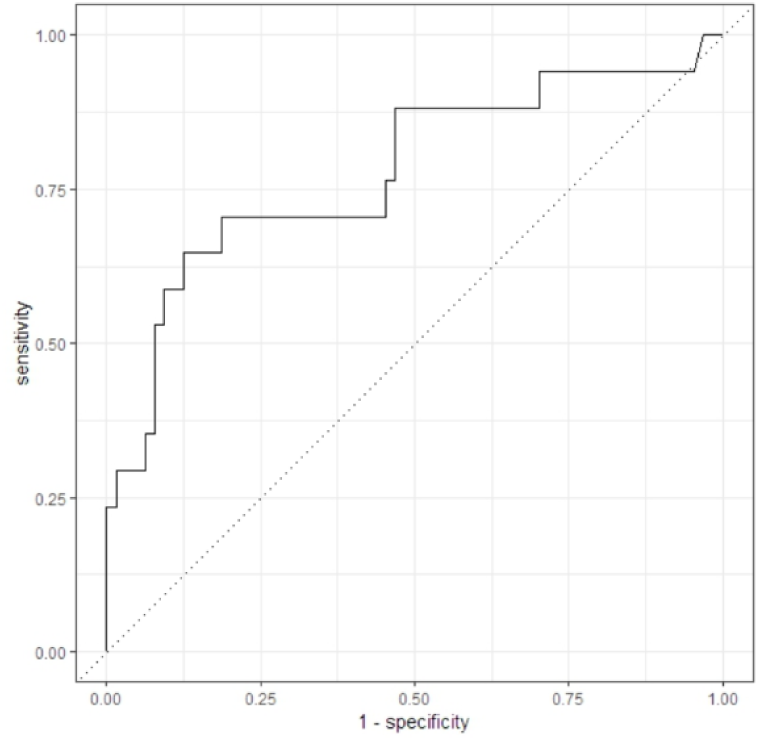
Receiver operating characteristic curve of the random forest model

**Table 4.** Performance metrics of the random forest model.

| accuracy | sensitivity | specificity | MCC | F<sub>1</sub>-score | AUC |
| --- | --- | --- | --- | --- | --- |
| 0.815 | 0.65 | 0.859 | 0.478 | 0.595 | 0.790 |

### Importance of the model variables

Figure 4 depicts the 10 most influential variables. MAPK9 (P45984) had the strongest influence on the classification model, and this DEP was significantly enriched in the IL-17 signaling pathway. Age, ADM (P35318), HTR7 (P34969), SLC44A2 (Q8IWA5), and GRM4 (Q14833) are among the most important variables, as expected. The remaining four variables are NTNG1 (Q9y2I2), FTSG3 (Q8IY81), CCAR2 (Q8N163), and UPS25 (Q9UHP3), in descending order of importance. In the poor cognition group, 7 of those 9 DEPs were upregulated, whereas SLC44A2 and GRM4 were downregulated (see Fig 5).

**Fig. 4.**
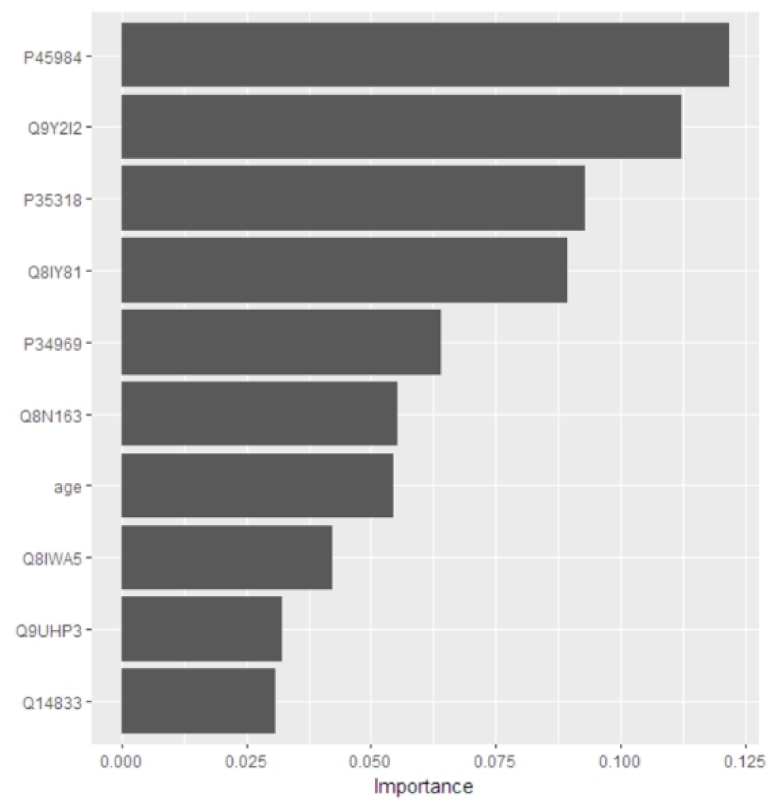
Significance of variables in the random forest model

**Fig. 5.**
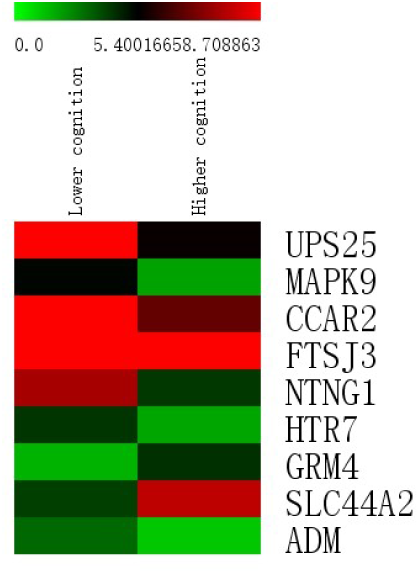
Heat map of the top 9 protein variables

## Discussion

This study built a classification model based on machine learning algorithms to detect healthy Thai subjects with poor cognitive performance from proteomic data. The random forest model showed good accuracy (0.815), specificity (0.859), and AUC (0.790), along with limited sensitivity (0.65). Overall, the screening performance of our classifier seemed to be acceptable [48, 49]. The performance of this model surpasses that of a previous study that utilized cardiovascular risk factors to compute the probability of mild cognitive impairment in the Thai population [50]. Despite this, due to its limited sensitivity and moderate F_1_-score [51], the performance of the random forest model should be considered cautiously. The low sensitivity was anticipated since the dataset was highly skewed toward the negative cases (higher cognitive ability) [52, 53] and the sample size was moderate [54, 55].

MAPK9 had the most impact on the random forest model; it was enriched in the IL-17 signaling pathway along with another major important variable, UPS25, and both DEPs were upregulated in the lower cognitive ability group samples. IL-17 signaling regulates inflammation by modulating inflammatory gene expression in cells [56]. These proinflammatory factors, if unrestrained, lead to the pathology of a variety of autoimmune and chronic inflammatory conditions [57]. Dysfunctional and persistent inflammatory processes were suggested as potential causative drivers of impaired cognitive functioning [58, 59]. This is consistent with previous findings that proinflammatory molecules can cause the progression of brain deficits [60], and IL-17 reportedly initiates the onset of synaptic and cognitive impairments in the early stages of Alzheimer’s disease [61], a process that may be mediated by activation of IL-17 receptors and MAPK [62].

Furthermore, serotonin receptor 7 (HTR7), metabotropic glutamate receptor-4 (GRM4), and choline transporter-like protein 2 (SLC44A2) were also among the most influential variables. It has been demonstrated that inhibiting HTR7 modulates immune responses and decreases the severity of intestinal inflammation [63]. Serotonin stimulation of HTR7 activates downstream signaling modules such as MEK/MAPK [64]. This suggested that HTR7 may be positively connected with IL-17-mediated neuroinflammation and the poor cognitive functioning resulting from brain inflammation. This notion is supported in part by a prior result that HTR7 antagonism may have positive effects on schizophrenia-like cognitive deficits [65].

According to previous animal research, GRM4 controls adaptive immunity and restrains neuroinflammation [66]. Another study found that mutant mice lacking GRM4 were less capable of learning and integrating new spatial information into previously generated memory traces [67]. The involvement of spatial information learning in the WCST process has been proposed [68]. Choline transporter-like protein 2 (SLC44A2) is a high-affinity choline carrier [69]. Choline is an indispensable constituent in the biosynthesis of acetylcholine (ACh), and its transportation into the presynaptic terminals of cholinergic neurons requires a high-affinity choline transporter [70]. ACh is assumed to play an essential role in executive function, and cholinergic decline is related to poor WCST performance in healthy individuals [71]. Besides, SLC44A2 is a newly discovered plasma membrane and mitochondrial ethanolamine transporter [72], and ethanolamine has been linked to anti-inflammatory effects [73]. The downregulation of GRM4 and SLC44A2 in the lower cognitive ability group samples provides a potential biological mechanism for why subjects in this group performed poorly on the WCST. However, the weights of SLC44A2 and GRM4 are small, consistent with there being multiple interactions across diverse neurotransmitter systems in maintaining central nervous system homeostasis [74, 75].

Netrin-G1 (NTNG1) and pro-adrenomedullin (ADM) are the second and third most important variables, and both DEPs were upregulated in the lower cognitive ability group samples. NTNG1 is a member of the Netrin family that is little studied in the brain but is implicated in inflammatory processes, including microglial function [76]. ADM has long been thought to be a biomarker for cognitive impairment [46, 77]. ADM accumulation in human brain contributes to memory loss with age [78], and blood ADM levels elevate in multiple pathological states, such as acute ischemic stroke and vascular cognitive decline with white matter alternations [79]. There is evidence indicating that inflammatory states stimulate the in vivo production of ADM [80]. These data, together with the evidence shown above, suggest that those in the lower cognitive ability group may have higher levels of neuroinflammation, resulting in poor cognitive performance on WCST. Age ranks as the seventh most important predictor in the model, with a moderate weight. Although it is widely accepted that aging is positively correlated to cognitive decline [33, 81], our findings imply that this variable did not emerge as an influential factor in classifying cognitive variability among healthy Thai subjects.

This study has limitations that should be noted. First, the lower and higher cognitive ability groups were defined by their WCST % Errors > 1SD from the mean, but there is a controversy about this cutoff for defining an abnormal score on the WCST [25], and further study with different cutoff points might be needed to replicate the findings of the current study. Second, only age and proteomics data were included in the random forest model. Given the potential direct and indirect interactions among multiple factors influencing cognitive function [7], additional factors such as socioeconomic status, lifestyle factors, and cardiovascular risk factors should be included in the model to improve model performance. Thirdly, while the WCST has been shown to be valid and reliable as a stand-alone cognitive assessment tool for healthy individuals of various ages and educational backgrounds [82], and %Errors offers a general measure of performance on the WCST that closely correlates with FSIQ [22], variability in different domains of cognition requires a more comprehensive battery of cognitive testing. This would have been beyond the scope of the current study. Last, due to the moderate sample size and imbalanced number of subjects in the lower and higher cognitive ability groups, generalizing the current findings should be done with caution.

## Conclusion

This study investigated the association between proteomics data and variability in normal cognition measured by the WCST. The results suggested that a measure of poor WCST performance in healthy Thai subjects might be attributed to higher levels of neuroinflammation. This study employed machine learning approach to shed light on the biological mechanisms that contribute to the variability in normal cognition.

## Acknowledgements

This work was partially supported by Global and Frontier Research University Fund, Naresuan University (grant number R2567C003) and Reinventing University Program 2024, the Ministry of Higher Education, Science, Research and Innovation (MHESI), Thailand (grant number R2567A141). The authors would like to thank all participants in this study. We would also like to thank the Faculty of Medical Science, Naresuan University and The National Center for Genetic Engineering and Biotechnology, Pathum Thani, Thailand for the facility supports.

